# Spatial IMIX: A Mixture Model Approach to Spatially Correlated Multi-Omics Data Integration

**DOI:** 10.1101/2023.07.15.549148

**Authors:** Ziqiao Wang, Bogdan Czerniak, Peng Wei

**Affiliations:** Department of Biostatistics, Bloomberg School of Public Health, Johns Hopkins University, Baltimore, MD, USA; Department of Pathology, The University of Texas MD Anderson Cancer Center, Houston, TX, USA; Department of Biostatistics, The University of Texas MD Anderson Cancer Center, Houston, TX, USA

## Abstract

Spatial high-throughput omics data allow scientists to study gene activity in a tissue sample and map where it occurs at the same time. This enables the possibility to investigate important early cancer-initiating events occur in normal-appearing tissue and gene activities that progress and carry through tumor tissue, as defined by “field effect.” The “field effect” genes are differentially expressed or methylated genes in the spatially resolved high-dimensional datasets with respect to the pathology subtype in each geographical sample across the tissue region. Current statistical methods for spatially resolved genomics data focus on the association of omics data with spatial coordinates without being able to incorporate and test for the association with the sample subtypes. In addition, analytical methods are underdeveloped for spatially resolved multi-omics data integration. We propose a novel statistical framework ‘spatial IMIX’ to integratively analyze spatially resolved high-dimensional multi-omics data associated with a specific trait, such as sample subtypes while modeling the spatial correlations between samples and the inter-data-type correlations between omics data simultaneously. Through extensive simulations, spatial IMIX demonstrated well-controlled type I error, great power by relaxing the independence assumptions between data types, model selection features, and the ability to control FDR across data types. Data applications to a geographically annotated tissue area of bladder cancer discovered cancer-initiating gene activities and revealed interesting fundamental biological mechanisms through pathway analysis. We have implemented our method in R package ‘spatialimix’ available at https://github.com/ziqiaow/spatialimix.

## 1 Introduction

High-throughput technologies have grown and evolved to accomodate new discoveries and insights in cancer research. Since studies often use multiple types of omics data, a wide variety of statistical methods have been developed over the past several years for integrative genomic analyses. Spatially resolved omics data add another layer, spatial information, into current multi-omics data-integration frameworks. In biology, spatial information allows scientists to investigate the complex interactions within and between biological networks (Wei and Pan, 2012; Zhu et al., 2007) in which each unit influences and is influenced by its neighboring environment. This is especially important for studies of cancer because tumors often contain complex mixtures of cell types in a single tissue area. Thus, incorporating structural and spatial information will help us better comprehend the dynamics of tumorigenesis, tumor development, and the tumor microenvironment.

Spatial high-throughput omics data allow scientists to investigate gene activity in a tissue sample and map where it occurs at the same time. For example, it is known that epithelial origin bladder cancers evolve from microscopically recognizable dysplasia. However, previous work (Czerniak et al., 2016; Majewski et al., 2019) showed important early cancer-initiating events occur in normal-appearing tissue, as defined by “field effect.” The genes involved in the development of early field effect control diverse cell functions and are expected to promote cell survival and proliferation, leading to the progression of neoplasia. Our motivating data example was based on geographically annotated mucosal samples from a surgically removed bladder specimen from one bladder cancer patient. Each spatial sample was evaluated microscopically and classified by a pathologist into one of three categories: normal urothelium (NU), *in situ* precursor lesions, or urothelial carcinoma (UC). The *in situ* precursor lesions were further dichotomized into low-grade intraurothelial neoplasia (LGIN) and high-grade intraurothelial neoplasia (HGIN). Furthermore, each spatial sample was measured for two whole genome-wide omics data platforms, gene expression and methylation. This study aimed to explore the cancer-initiating events that occur in normal-appearing tissue samples that carries on to carcinoma samples in a single tissue section, i.e., discover differentially expressed and methylated genes in the spatially resolved high-dimensional datasets with respect to the sample subtypes across the tissue, and furthermore the fundamental biological mechanisms. Another example is the spatial transcriptomics, which profiles gene expression in its spatial context in tissues, allowing scientists to investigate the complex mix of cell types and structures in tissues, including tumors and their micro-environments (Berglund et al., 2018).

Current statistical methods for analyzing high-dimensional spatially correlated data mostly aim to identify and characterize spatially variable genes, such as SpatialDE (Svensson et al., 2018) and SPARK (Sun et al., 2020). Both methods build upon a generalized linear spatial model with different link functions targeting at respectively normalized expression data and count data. To test the hypothesis whether a gene shows spatial expression variance across the tissue area, both methods test the random effect term in the spatial mixed model. Different groups of spatial variance pattern of the identified genes are then determined by an unsupervised clustering method, such as hierarchical clustering, which is incapable of incorporating a specific sample location’s subtype into the analysis. Thus, these methods cannot address the bladder cancer whole-organ mapping data problem as described above, which involves spatial sample subtypes (NU, LGIN, HGIN, and UC) across the whole bladder tissue and identification of genes that are differentially expressed or methylated in the same direction across all spatial locations with nonspatial variability. Furthermore, given that spatially resolved proteomics, epigenomics, and metabolomics technologies are still under development, there is as yet no statistical method for multi-omic spatial data integration, to our knowledge.

Therefore, we developed spatial IMIX, a method for identifying genes in association with covariates, such as sample subtypes, through multiple spatially correlated omic data types (Fig. 1). This method focuses on the fixed effect of sample subtypes in a spatial linear mixed model that incorporates the spatial correlations on a two-dimensional surface. For data integration, it models the summary statistics *z*-scores, which are transformed from the *P* - values of the fixed effect estimates, with a multivariate Gaussian mixture distribution. This step additionally characterizes the correlations between various data sources. This method extends the previous work by Wang and Wei (2020), to incorporate spatial correlations between samples. Our method is computationally efficient because it uses the expectation-maximization (EM) algorithm for parameter estimation. It also controls the false discovery rate (FDR) and features statistically principled model selection. We have implemented our method in R package ‘spatialimix’ available at https://github.com/ziqiaow/spatialimix, as well as CRAN soon.

**Figure 1:**
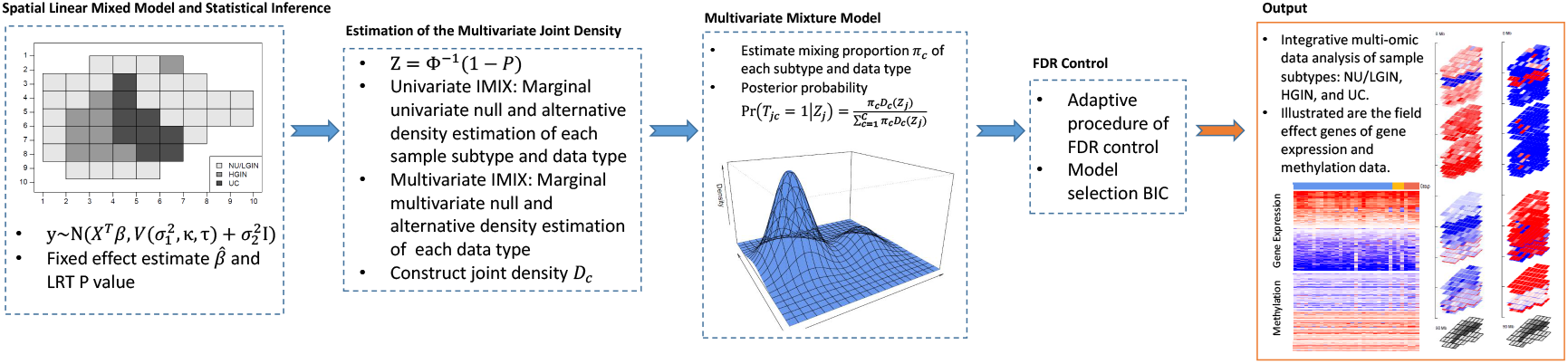
Spatial IMIX model framework.

## 2 Methods

### 2.1 Spatial Mixed Model and IMIX for One Data Type

#### Spatial Mixed Model

We first consider modeling one type of spatially resolved omics data, such as gene expression or methylation data. Each data type is measured for *p* genes and *n* spatial locations/samples on a tissue of interest. Our motivating example uses geographically annotated mucosal samples from a surgically removed bladder specimen from one bladder cancer patient. Here, the omics data in our motivating example are measured on the whole genome, with the number of genes *p* approximately equal to 20 000. Each sample *i* (*i* = 1, …, *n*) is spatially correlated over a two-dimensional space. The spatial coordinate of each sample is *s*_*i*_ = (*s*_*i*1_, *s*_*i*2_) ∈ ℝ^2^. Note that *s*_*i*_ could be extended to more than two dimensions, for instance for tissues with three-dimensional coordinates such as height, width, and length or even a fourth-dimension such as the distance between different time points (Sun et al., 2020). Consider a categorical variable that contains three classes: NU/LGIN (LG), HGIN (HG), and UC. Each sample *i* belongs to one of the three classes. Here, we want to identify the genes that show field effect (i.e., genes that are differentially expressed/methylated in all three classes, LG, HG, and UC, compared with the controls); genes that are differentially expressed/methylated in HG and UC samples; and genes that are only differentially expressed/methylated in UC samples. The biological rationale for this was introduced in Section 1. We model the data using a spatial linear mixed model:

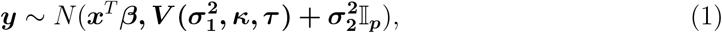

the matrix form can be expanded as:

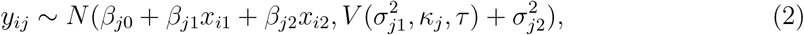

where *y*_*ij*_ is the log2 ratio of the expression value or methylation value of gene *j* in sample *i* (*i* = 1, *…, n*; *j* = 1, *…, p*) compared with the controls. *x*_*i*1_ is the indicator (dummy variable) of whether sample *i* is in the class HG, and *x*_*i*2_ is the indicator of whether sample *i* in the class UC. The indicators remain the same for any gene *j*. The fixed effect *β*_*j*0_ gives the overall mean for gene *j, β*_*j*1_ is the fixed effect of HG relative to LG, and *β*_*j*2_ is the fixed effect of UC relative to LG. 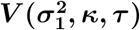 is the spatial covariance matrix defined on the distance between each pair of samples/locations and 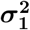 is a vector of variance components (Li et al., 2009). Here, for gene *j*, the (*i, i*)th component 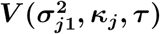 is 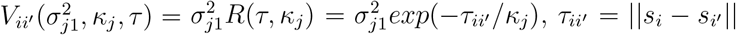, *τ*_*ii*_ = ||*s*_*i*_ − *s*_*i*_ || is the Euclidean distance between any two samples *i* and *i* for any gene *j*, and *κ*_*j*_ is a parameter that controls how fast the correlation decays with distance, note that *κ*_*j*_ *>* 0. A larger *κ* indicates a stronger correlation between two samples, and therefore smaller semi-variance, which is defined as 1 − *exp*(−*τ/κ*). The exponential spatial structure is a special case of the Matern correlation structure 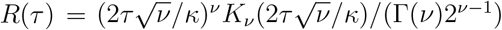 when 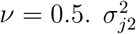 is the independent non-spatial error.

#### Statistical Inference

To identify differentially expressed/methylated genes in a certain combination of groups (for example, the field effect, LG+,HG+,UC+), we perform hypothesis testing for each group *g, g* = 0, 1, 2. The null hypothesis of each fixed effect estimate *β*_*jg*_ can be formulated as 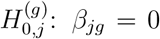, i.e., gene *j* is not differentially methylated/expressed in group *g*. For statistical inference, we use likelihood ratio tests (LRTs) based on the maximum likelihood estimation of the spatial mixed model. We compare the model likelihood of the fitted spatial mixed model to the likelihood of the null model. For example, to test 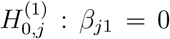, i.e., gene *j* is not differentially methylated/expressed in HG, we compare the full model with the null model 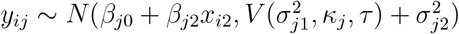. Genes that are significantly differentially expressed/methylated between groups are commonly identified by applying the Benjamini-Hochberg FDR method to adjust for multiple testing in each group and then simply determining the overlap of significant genes between groups. However, this strategy may reduce statistical power because it assumes that the fixed effect estimates are independent of each other and omits other unknown dependence structures apart from the spatial correlations of each group. In addition, the FDR control is performed for each group separately without considering the across-group FDR.

#### Extension of IMIX

Therefore, we propose a new method to address the above problems. We take the summary statistics *P* -value *p*_*jg*_ from the LRT results fitted by the spatial mixed model, and use the multivariate mixture model approach IMIX developed by Wang and Wei (2020) to test for differentially expressed/methylated genes in a certain combination of groups. The *P* -values are transformed to *z*-scores *z*_*jg*_ by *z*_*jg*_ = Φ^−1^(1 − *p*_*jg*_), where Φ is the cumulative distribution function of the standard normal distribution *N* (0, 1) (McLachlan et al., 2006; Wei and Pan, 2008). To simplify the method, we only applied IMIX-Ind and IMIX-Cor-Twostep on the *z*-scores. Here, we assume that each group combination corresponds to a latent state. Table 1 shows the group combinations of interest and their corresponding latent state components in IMIX. Possible IMIX components by order are (0, 0, 0), (1, 0, 0), (0, 1, 0), (0, 0, 1), (1, 1, 0), (1, 0, 1), (0, 1, 1), (1, 1, 1).

**Table 1:**
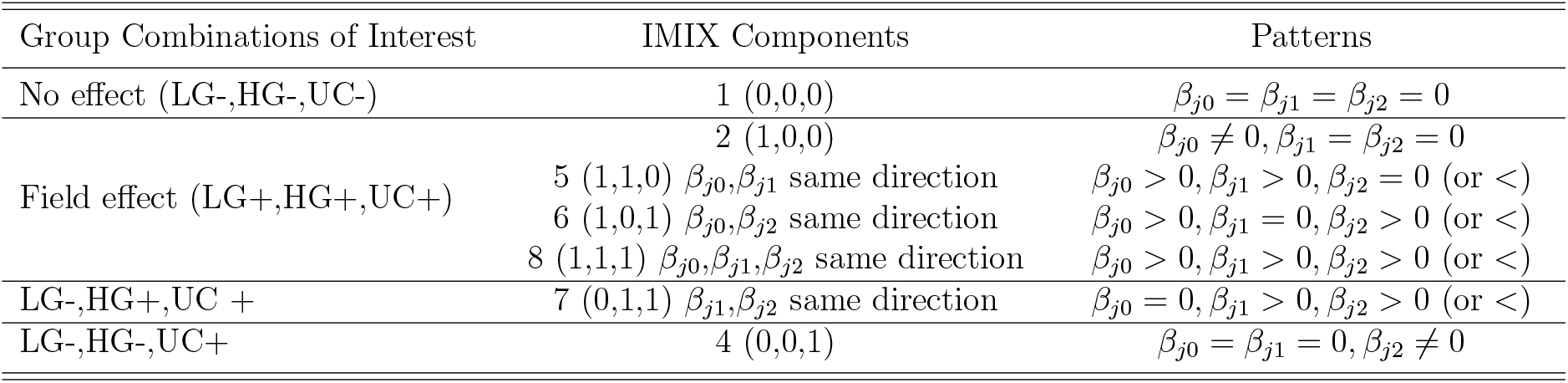
Patterns of fixed effects corresponding to field effects in NU/LGIN (LG), HGIN (HG), and UC groups and IMIX components for fixed gene *j*. + represents a gene that is differentially expressed/methylated, - represents a gene that is not differentially expressed/methylated compared with the controls.

Other possible ways to perform the hypothesis testings are using the spatial mixed model with the F test (Pinheiro et al., 2021) for inference and using linear regression without considering the spatial correlations. We will further explore these alternatives in the Results section where we conducted simulation studies to evaluate the type I error and power.

### 2.2 Integrative Spatial IMIX for More Than One Data Type

In our motivating data example (section 3.2), there were two data types: gene expression and methylation data. The goal was to identify genes that showed field effect in both data types. A straightforward way to accomplish this is to test each data type separately using the methods described in section 2.1 and find the overlap of the significant genes in both data types. We propose two integrative models to directly analyze both data types simultaneously by accounting for the underlying dependence structures between multiple data types. We call these two methods univariate spatial IMIX and multivariate spatial IMIX. A special feature of these methods is that samples of multiple data types can be overlapping, different, or the same because the models use the summary statistics for each data type. Without loss of generality, this section focuses on *H* = 2 data types. The main idea is that we assume the summary statistics *z*-scores of the genes, which are retrieved and transformed from the spatial mixed model fitting *P* -values, follow a multivariate Gaussian mixture model. Each component corresponds to a group combination, for example, component 1 corresponds to (LG-_1_,HG-_1_,UC-_1_,LG-_2_,HG-_2_,UC-_2_). Here 1, 2 are data types 1, 2, respectively, i.e., the genes in component 1 in the mixture model are not differentially expressed/methylated in any of the disease classes for either data type. In total, there are *C* = 2^6^ = 64 components in the mixture model, corresponding to all possible group combinations. We estimate the probability density function *D*_*c*_, *c* = 1, *…*, 64 of each component and the proportion *π*_*c*_ of each component in the mixture model. In the statistical formula, we assume that *Z*_*j*_ = (*z*_*j*11_, *z*_*j*12_, *z*_*j*13_, *z*_*j*21_, *z*_*j*22_, *z*_*j*23_)^*T*^ comes from a mixture distribution with *C* = 64 mixture components:

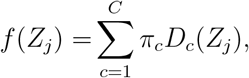

where each component *c* follows a six-dimensional multivariate distribution *D*_*c*_, and the mixing proportions are *π*_*c*_, *c* = 1, *…*, 64, subject to 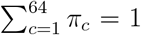. Depending on the latent state of gene *j*, i.e., whether it belongs to latent state *c* or not, we have *T*_*jc*_ = 1 or *T*_*jc*_ = 0, respectively. To assess how likely gene *j* belongs to the latent state/component *c*, we estimate the posterior probability of the latent label *T*_*jc*_:

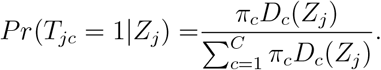

We further assume the *c*-th component distribution *D*_*c*_ to follow a multivariate Gaussian distribution (Wang and Wei, 2020). The marginal mixture density *f* (*Z*_*j*_) can then be written as 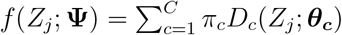, where *D*_*c*_(*Z*_*j*_; ***θ***_***c***_) = *ϕ*(*Z*_*j*_; ***µ***_***c***_, **Σ**_***c***_). In the following sections we will describe how we estimate the unknown parameters of the marginal density functions. Then we use the EM algorithm (Dempster et al., 1977) to estimate the mixing proportions ***π*** of the latent states/components *C*.

#### 2.2.1 Estimation of the Multivariate Joint Density

##### Univariate Spatial IMIX

For *H* = 2 data types and *G* = 3 groups of interest (LGIN, HGIN, UC) within each data type, we estimate the marginal empirical null and the alternative densities first. The null hypothesis is 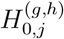: gene *j* is not differentially methylated/expressed in group *g* of data type *h*. Since the groups in each data type are independent of each other, and that the data types are independent of each other, we assume

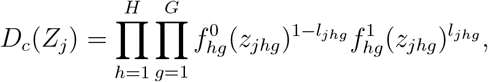

The complete data vector is

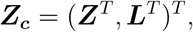

where

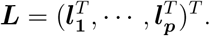

The label vectors ***l***_***j***_ = (*l*_*j*11_, *…, l*_*jHG*_), *j* = 1, *…, p* are binary variables used to denote each latent state *c* of gene *j*. If *l*_*jhg*_ = 1, gene *j* is differentially expressed/methylated in group *g* of data type *h*; and vice versa. For example, if ***l***_***j***_ = (1, 1, 1, 0, 0, 0), the joint density *D*_*c*_ of gene *j* is modeled as the product of the alternative marginal density functions from all three groups in data type 1 and the null marginal density functions from all three groups in data type 2: 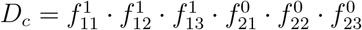.

Intuitively, we assume that the *z*-scores of genes in group *g* of data type *h*, as described previously in Section 2.1, follow a univariate Gaussian mixture model (Wei and Pan, 2008):

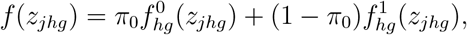

here *z*_*jhg*_ is the *z*-score of gene *j* in group *g* of data type *h*, 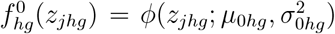 and 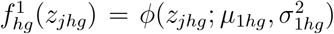 are the probability density functions following a normal distribution. In particular, *µ*_0*hg*_ and 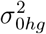 correspond to the null hypothesis, and *µ*_1*hg*_ and 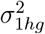 are the alternatives in group *g* of data type *h*. Note that the *z*-score transformation ensures that smaller *P* -values are transformed to larger *z*-scores, which correspond to the alternative hypothesis, i.e., that the distribution of the *z*-scores under the alternative hypothesis (alternative distribution) has a larger mean than does the null distribution (McLachlan et al., 2006). Therefore, the mixture components for the null and the alternative hypotheses can be easily distinguished. We use the EM algorithm to estimate the parameters and construct multivariate joint density functions with the estimated mean 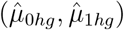 and variances (*vâr*_0*hg*_,*vâr*_1*hg*_), *h* = 1, 2, 3; *g* = 1, 2. Therefore, the constructed density function *D*_*c*_ is a six-dimensional Gaussian distribution here. Based on Gleason et al. (2020), we approximate the density function of each component *c* to be *D*_*c*_ ∼ *N* (***µ***_***c***_, **Σ**_***c***_) with the mean vector as 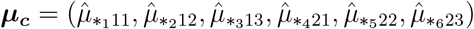 and the diagonal variance-covariance matrix as 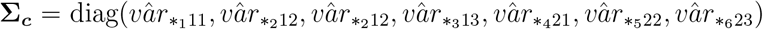. Here, *_1_, *_2_, *…*, *_6_ can be either 0 or 1 and they are coherent between the mean vectors and the variance matrices. The univariate spatial IMIX method is flexible and can be easily extended to more groups and more than two data types.

##### Multivariate Spatial IMIX

To estimate the unknown parameters ***µ***_***c***_, **Σ**_***c***_ of the multi-variate density functions *D*_*c*_, we first fit the spatial mixed model for each data type separately and use the IMIX extension as described in Section 2.1. Similarly to the univariate spatial IMIX, the estimated parameters in the first step are used to approximate the parameters in the joint model with new mean vectors and new variance-covariance matrices as block diagonal matrices using the estimated variance-covariance matrices for each data type. This model is less parsimonious than the univariate spatial IMIX method, although the simulation study will prove that both methods are effective.

#### 2.2.2 Adaptive Procedure for FDR Control

For each component *c* (*c* ≠1), we construct the following hypotheses:

- 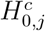: Gene *j* does not belong to component *c*;
- 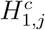: Gene *j* belongs to component *c*.

Note that component 1 (the global null) is not considered as a “discovery” for which FDR is applicable. The estimated posterior probability that gene *j* belongs to component *c* is defined as 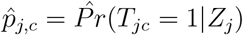 and the estimated local FDR for gene *j* is defined as 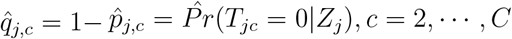. We adapt the across-data-type FDR control procedure in the IMIX framework (Sun and Cai, 2007; Wang and Wei, 2020) for the integrative spatial IMIX models: let 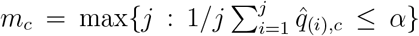, we reject all *H*_0,(*j*)_, *j* = 1, *…, m*_*c*_, where 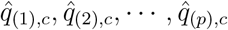 are ranked in component *c*. This adaptive procedure controls the FDR for each component at level *α* asymptotically under weak conditions (Genovese and Wasserman, 2002). We can extend this procedure to control the FDR under more than one component. For instance, if we want to identify the genes that show field effect in one data type, the procedure can be applied to the combination of components 2, 5, 6 and 8 as illustrated in Table 1.

## 3 Results

### 3.1 Simulation Study

We performed three sets of simulation studies. The first set was to evaluate the type I error and power of the spatial mixed model, the second set of simulations was to investigate our method for one data type and compared them with linear regression, spatial mixed models using the F test and LRT for fixed effect inference, and the second set was for integrative data analysis of two data types. In addition, we evaluated the AIC and BIC model selection of the spatial integrative models using the third set of simulated data.

#### 3.1.1 Type I Error and Power Investigation

We assessed the type I error and power for fixed effect testings using spatial mixed model. We simulated 10 000 datasets using 45 samples (*j* = 1) mimicking the spatial locations in the bladder cancer whole-organ mapping data. There were 27 LG, 9 HG, and 9 UC samples in the simulated data. Here, the dataset followed a spatial linear mixed model with exponential correlation structure. We tested the type I error and power at *α* = 0.05. For power simulations, we simulated the three fixed effects respectively follow normal distribution with standard deviation equals to 0.1 and mean as shown in Table S2 & S4. In each scenario, all three fixed effects followed the same distribution. We compared the joint testing of three fixed effects using linear regression based on F test and Wald test, the spatial mixed model based on Wald test and the spatial mixed model with LRT (smm+LRT). For the hypothesis testing for each fixed effect, we compared linear regression, the spatial mixed model with the default F test for fixed effect statistical significance (smm+Ftest), and smm+LRT. More details are in the supplementary materials Section 1.1. The results of joint testing (Table S1) showed that only smm+LRT could control type I error close to 0.05, while linear regression and mixed model using the Wald test failed to control the type I error under various spatial variance settings. The separate testing (Table S2) showed that smm+Ftest and smm+LRT could control type I error for HG and UC when sample sizes were small, however, smm+Ftest failed to control the type I error for LG group. The power of smm+LRT, which controlled type I error well under all scenarios, was close to the power of those methods that failed to control the type I error (Table S1 & S2). Therefore, we adopted the *P* values of smm+LRT for further IMIX input in this study.

#### 3.1.2 Simulation for One Type of Data

We simulated data *y*_*ij*_, *i* = 1, *…*, 45, *j* = 1, *…*, 1 000 based on equation 2. Here, the spatial coordinates and the groups of interest were based on the bladder cancer whole-organ mapping data described in Section 3.2. We replaced the missing samples from the real data set with HG and UC groups. In total, the simulated data contained 45 spatial locations, with 27 spatial samples in the LG group, 9 samples in the HG group, and 9 samples in the UC group, and 1 000 genes. We simulated four scenarios of 1 000 datasets each to evaluate the methods for the spatial random effect variances 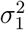, they are (0.1, 0.5, 1, 2), respectively. The spatial correlation followed an exponential structure. The independent non-spatial residual 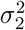 was set as 1. We considered four groups of fixed effects *β*_*j*0_, *β*_*j*1_, *β*_*j*2_ following a multivariate Gaussian distribution with standard deviation 0.1 and correlation matrix as identity matrix across all groups and mean vectors (0,0,0), (2,0,0), (0,2,2), (0,0,2) for 250 genes respectively. Here each group corresponds to genes in the no effect, field effect, HG & UC+, and UC+ only groups.

We applied our proposed spatial IMIX method, the spatial mixed model with IMIX extension (smm+IMIX), smm+Ftest, smm+LRT, and linear regression. We set *α* = 0.2 as the nominal error control level across all methods for comparisons. Here, smm+IMIX controlled FDR across all groups. smm+Ftest, smm+LRT, and linear regression used the Benjamini-Hochberg FDR at 0.2 for *β*_*j*0_, *j* = 1, *…*, 1 000, *β*_*j*1_, *j* = 1, *…*, 1 000, *β*_*j*2_, *j* = 1, *…*, 1 000 respectively, to infer the statistical significance. The groups were assigned based on the patterns shown in Table 1. Note that for smm+IMIX, we first fitted the spatial mixed model and used LRT to extract the summary statistics *P* -values as the input for IMIX testing. LRT was able to control the type I error well in the spatial correlation settings as described in Section 3.1.1.

Figure 2 presents the simulation results for the average of 1 000 simulations of the misclassification rate, with five class labels: no effect, field effect, HG & UC+, UC+ only, and others. As the spatial random effect variance increased, misclassification rates increased for all models. In particular, linear regression had the highest misclassification rate and standard deviation among all methods because it did not model the spatial correlations within the data. smm+LRT performed slightly better than smm+Ftest when the spatial random effect variance was relatively small. Note that in the last scenario, when 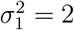, smm+LRT had a high average misclassification rate, resulting in a slightly increased standard deviation for the misclassification rates of smm+IMIX. However, overall smm+IMIX consistently achieved a lower misclassification rate in all scenarios than did the other methods.

**Figure 2:**
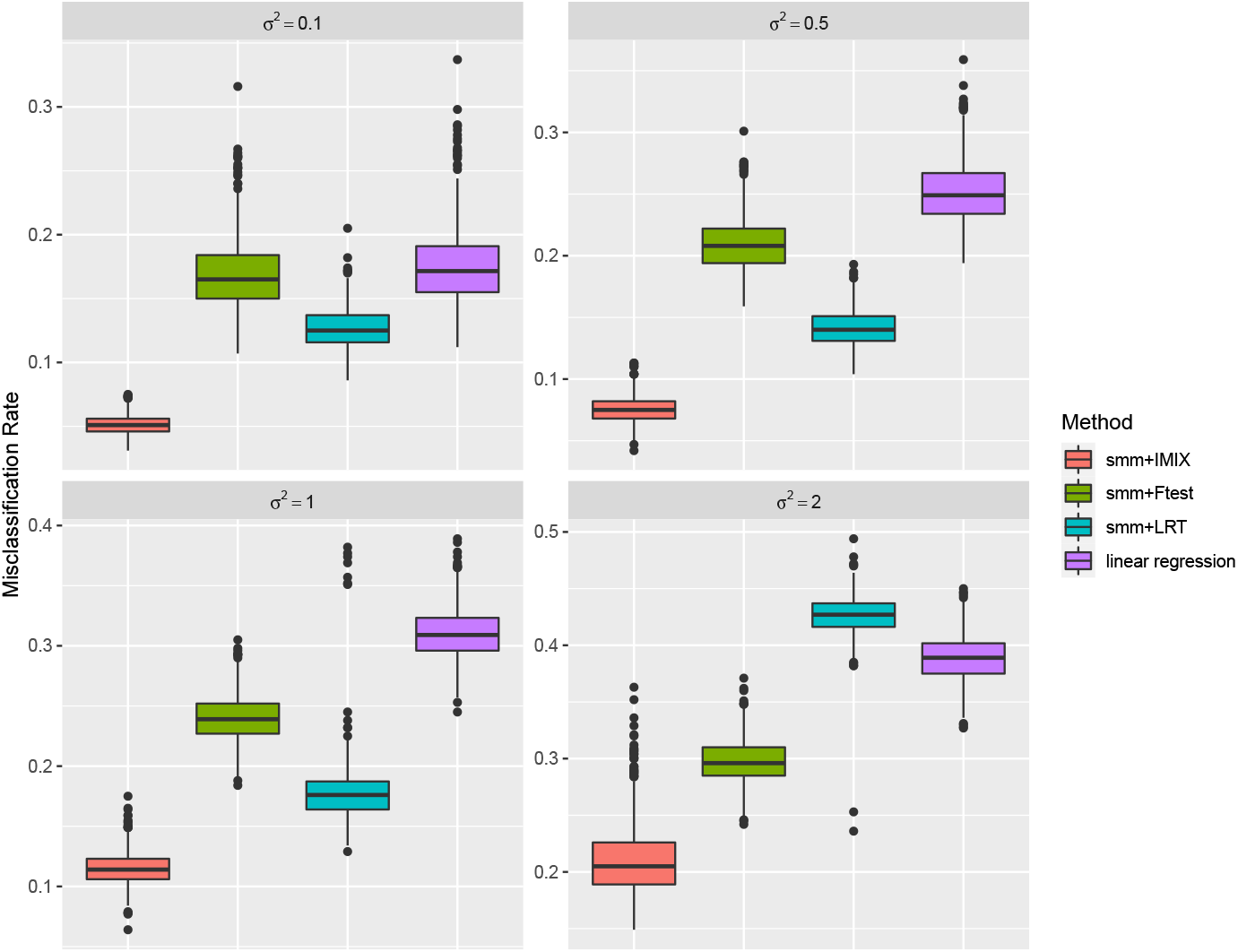
Simulation study results for one data type comparing spatial mixed model (smm) with IMIX extension, smm with F test, smm with likelihood ratio test (LRT), and linear regression using different spatial correlation variance settings.

#### 3.1.3 Simulation for Integrative Data Analysis

The second set of data *y*_*ijh*_ was simulated based on

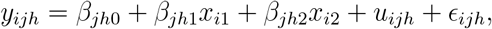

where *y*_*ijh*_ is the log2 ratio of gene *j* at location *i* of data type *h* (*i* = 1, *…, n*; *j* = 1, *…, p*; *h* = 1, 2) compared with the control. We fixed the samples to be the same across both data types. We fixed *β*_*j*10_ = *β*_*j*20_, *β*_*j*11_ = *β*_*j*21_ and *β*_*j*12_ = *β*_*j*22_ across the data types for field effect and monotonic changes. *u*_*ijh*_ is the spatial random effect for gene *j* in sample *i* of data type *h*. We set the spatial correlation following an exponential structure; note that *cor*(*u*_*ij*1_, *u*_*ij*2_) = 0, i.e., for a fixed gene *j* of sample *i*, the two data types are not spatially correlated. Here, (*u*_1*jh*_, *…, u*_*njh*_) is 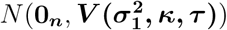, accounting for the variation within each data type. 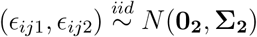 is a multivariate Gaussian error vector and independent of *u*_*ijh*_. Here, _*ijh*_ only influences the variation between two data types and does not account for the variation within each data type.

We simulated four scenarios of 1 000 datasets to compare the two integrative models and the separate IMIX models. We simulated varied effect sizes and residual covariance matrices **Σ**_**2**_ in the following scenarios. Scenario 1 assumed no correlation between two data types. In scenario 2, the independent error term of the two data types followed a multivariate normal distribution with 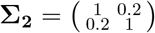 across the no effect, field effect, HG & UC+, and UC+ only groups. Scenario 3 followed the same settings as scenario 2 except we set a high correlation of 0.8. Scenario 4 was based on the real data in section 3.2 with correlations of 0.08763, 0.04206, 0.3, and 0.6814 between two data types in the four groups, respectively, and the standard deviation fixed at 1. In all scenarios, the dataset for each data type contained 1 000 genes with fixed spatial random effect variance 0.1 and an exponential correlation spatial structure based on the distance of 45 spatial locations; fixed effects *β*_*j*0_, *β*_*j*1_, *β*_*j*2_ followed multivariate Gaussian distribution with standard deviation 0.1 and correlation matrix as identity matrix across all groups. We set the mean vectors as (0,0,0), (*µ*\*,0,0), (0,*µ*\*,*µ*\*), (0,0,*µ*\*) for 250 genes respectively. Here, each group corresponds to genes with no effect, field effect, HG & UC+, and UC+ only. We set *µ*\* as 1 (small effect), 2 (medium effect), and 3 (large effect) respectively.

Again, we applied our proposed method, multivariate spatial IMIX, univariate spatial IMIX, and the methods we used for one data type, smm+IMIX, smm+Ftest, smm+LRT, and linear regression. We analyzed each data type separately and combined the results of two data types in an *ad hoc* manner. We set *α* = 0.2 as the nominal error control level across all methods for comparisons. Here, smm+IMIX, smm+Ftest, smm+LRT, and linear regression controlled the FDR for each data type separately; whereas multivariate spatial IMIX and univariate spatial IMIX controlled the FDR for both data types simultaneously. In particular, the multivariate spatial IMIX model used the estimates from the IMIX-Cor-Twostep model described in Wang and Wei (2020) for both data types.

Figure 3 and Figure S1 shows the simulation results for the average of 1 000 simulations of the misclassification rate, again with five class labels: no effect, field effect, HG & UC+, UC+ only, and others. As the effect sizes increased, all models achieved lower misclassification rates. Among all methods, smm+IMIX, multivariate IMIX, and univariate IMIX had the lowest misclassification rates. We did not observe substantial changes in the mis-classification rates within each method when the data correlation increased, owing to the noise imposed by the spatial correlation variations; however, we still observed significantly lower misclassification rates with IMIX methods compared with the other three methods. In particular, the two proposed integrative methods, univariate IMIX and multivariate IMIX, performed better compared with separate smm+IMIX under different correlation and effect size settings. In particular, univariate IMIX performed better than multivariate IMIX when the effect size was relatively large. Univariate IMIX is more parsimonious than multivariate IMIX, and in turn holds the advantage of better classification due to the bias-variance tradeoff. When the effect size was small, multivariate IMIX was slightly more robust than the other two methods.

**Figure 3:**
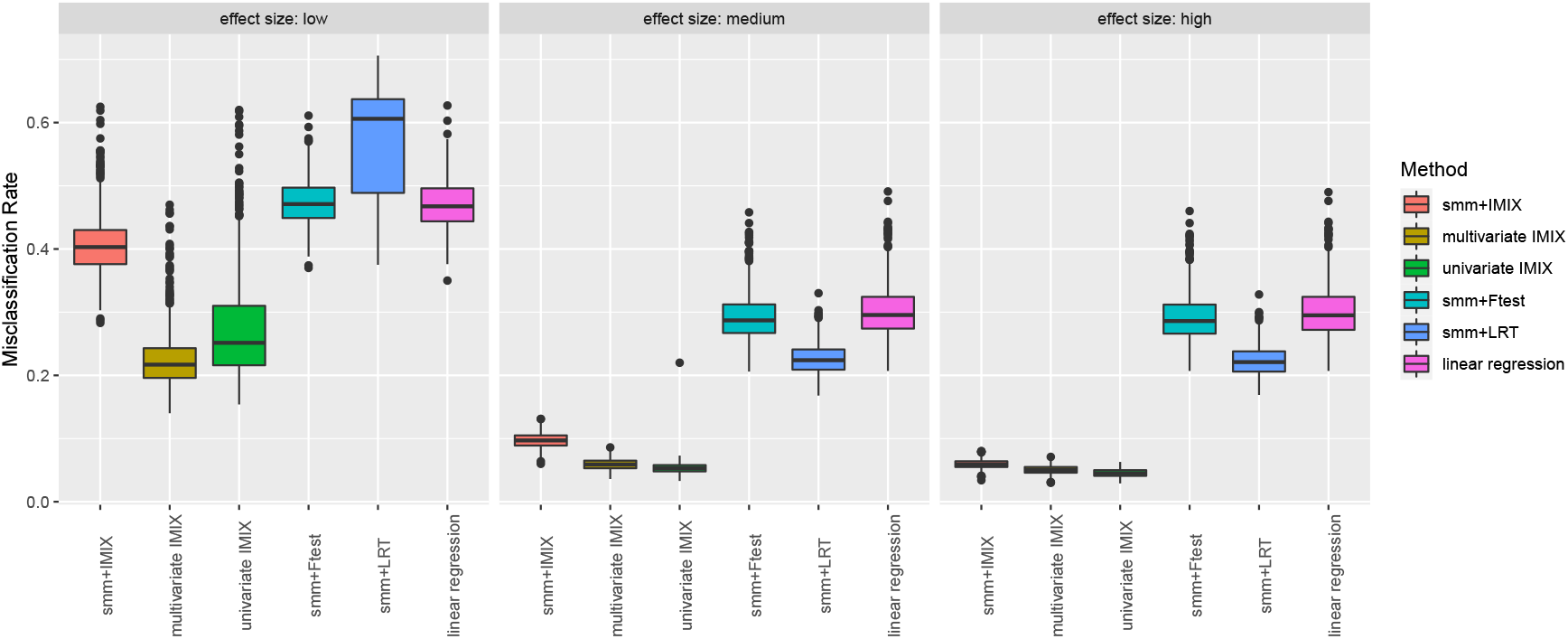
Simulation study results for two omics data types comparing spatial mixed model (smm) with IMIX extension, multivariate spatial IMIX, univariate spatial IMIX, smm with F test, smm with likelihood ratio test (LRT), and linear regression using different effect size settings with between-data-type correlations mimicking the bladder cancer whole-organ mapping data example (Scenario 4).

#### 3.1.4 Model Selection

We evaluated the performance of AIC and BIC for selecting the correct number of components in the integrative spatial IMIX framework. Here, we used the 1 000 simulated datasets described in section 3.1.3 with different effect size settings ranging from low to medium to high of scenario 4. The true underlying number of components was four across all three scenarios: no effect, field effect, HG&UC+, and UC+ only for both data types with balanced mixing proportions. There were in total 64 possible components in the mixture model, and our previous model fitting for simulation (see Section 3.1.3) was directly based on all components without considering the model selection. In this section, we investigate the model selection criteria and see whether they could further improve our model fitting and thus the misclassification rate.

Figure 4 shows the proportion of selected components in each simulation study based on AIC or BIC. Both model selection criteria were able to identify the true number of components in the simulated data. However, AIC performed worse than BIC when the effect size was small, especially for the univariate IMIX model. In addition, AIC tended to overestimate the number of components in both the multivariate and univariate IMIX models. BIC was more robust under different scenarios and models. Figure S2 shows the AIC and BIC values of all possible components after averaging 1 000 simulation replications for different effect sizes. Both AIC and BIC performed well in identifying the true number of components. We consider BIC to be more stable as it additionally considers the number of genes in the penalty term, which can be as large as tens of thousands under the whole-genome setting. We compared the misclassification rate between the four-component spatial IMIX models after model selection based on BIC values and the spatial IMIX models before model selection as described in section 3.1.3. Table S5 shows that the 4-component spatial IMIX models after model selection based on BIC performed better than the 64-component spatial IMIX model, especially when the fixed effect size was small. We conclude that model selection improves model fitting for spatial IMIX and results in more accurate classifications.

**Figure 4:**
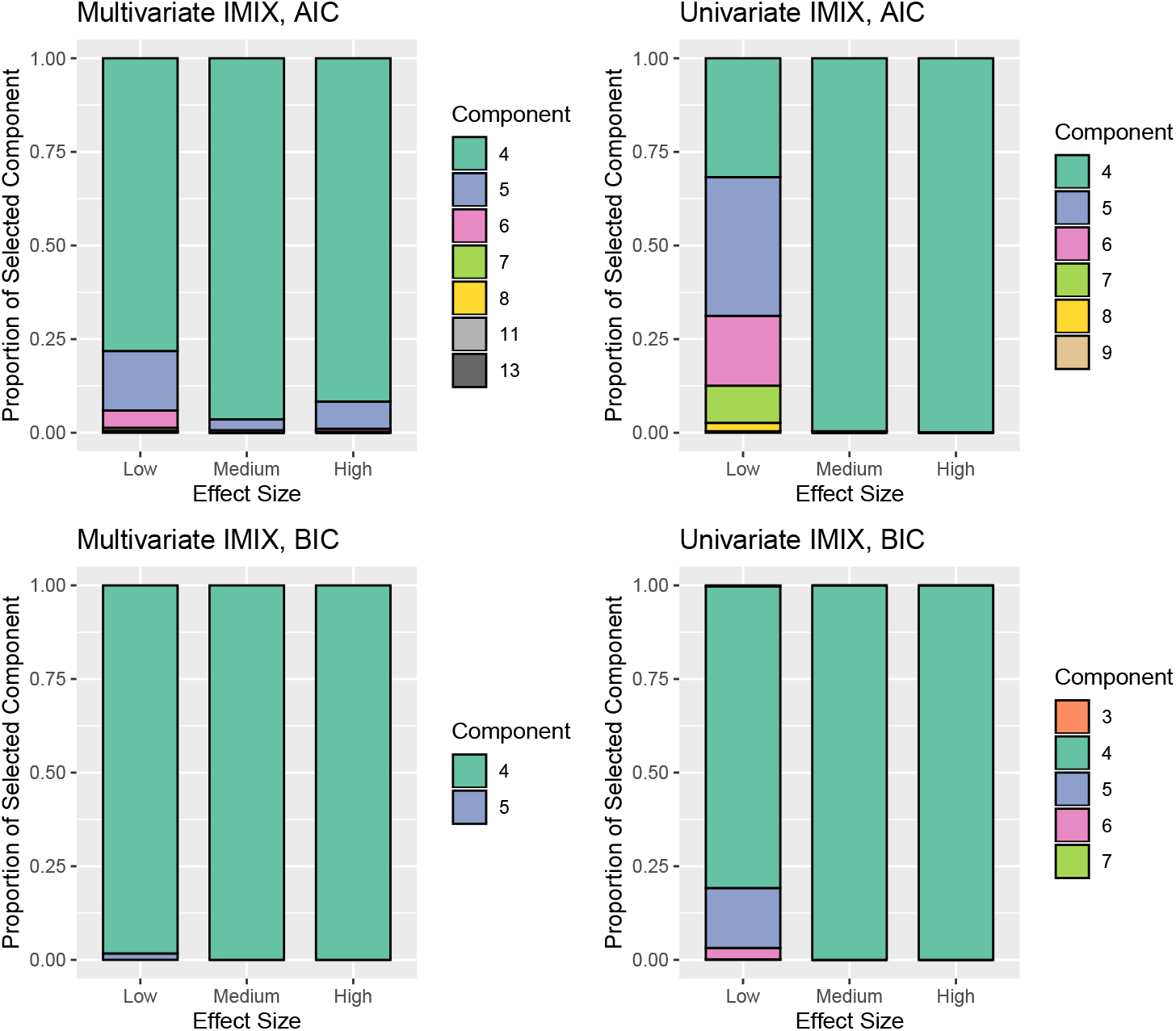
Proportions of selected components based on AIC and BIC of 1 000 simulation studies with different effect size settings.

### 3.2 Multi-Platform Whole-Organ-Based Molecular Profiling of Bladder Cancer

As described in Section 1, here we investigated cancer-initiating events in normal-appearing tissue that carry through to carcinoma, i.e., field effect, in epithelial basal-subtype bladder cancer. The data were obtained from geographically annotated mucosal samples collected from a surgically removed bladder specimen and further details can be found in Bondaruk et al., 2022. Each spatial sample was classified into three subtypes: NU/LGIN, HGIN, and UC. Each sample contained a two-dimensional spatial coordinate measured on two whole-genome-wide omics data platforms, gene expression and methylation. We aimed to identify genes associated with field effect in both data types across all disease subtypes and explore whether these genes reveal any interesting biological mechanisms.

The disease grades in individual samples of the cystectomy specimen classified as NU/LGIN, HGIN, and UC are shown in Figure 5(a). Together there were 35 spatial locations for both RNAseq and methylation data in the tumor specimen. Here, RNAseq and methylation data both were measured for 34 samples out of the spatial locations. For both data types, three HGIN and four UC samples were measured in the same locations. Each data types measured 27 NU/LGIN samples with 26 same locations and only one location that is different. There were eight control samples of methylation data and five control samples of gene expression data. The controls samples were measured from sex-matched normal urothelial suspensions, which were prepared from the ureters of nephrectomy specimens that were free of urothelial neoplasia (Majewski et al., 2019). The data preprocessing and normalization procedures are described in the supplementary materials Section 2.1. In total we had 18 604 genes in the gene expression data and 14 682 genes in the methylation data for downstream analysis with the integrative spatial IMIX models. We used the log2ratio between each sample in the disease group compared with the average of controls in the downstream analysis.

**Figure 5:**
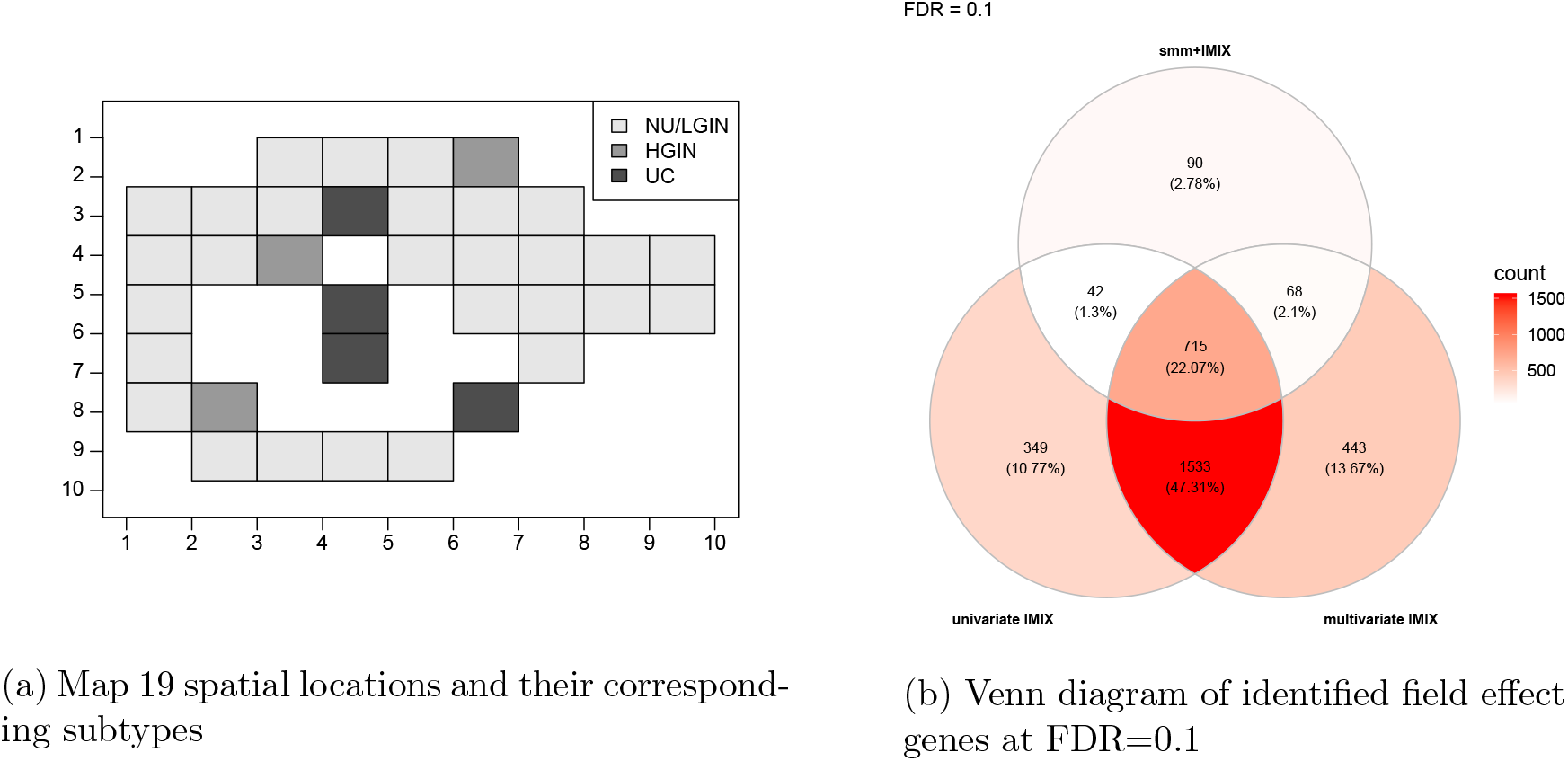
Whole-organ mapping data application visualizations. (a) The spatial locations of the disease grades of the whole organ mapping samples of map19 bladder cancer. (b) Comparisons of genes identified by smm+IMIX, univariate IMIX, and multivariate IMIX, FDR=0.1.

We applied our proposed spatial integrative data methods—univariate IMIX, multivariate IMIX, and smm+IMIX—on the spatially correlated high-dimensional data as described above. The results are summarized in Figure 5(b); the FDR was controlled at *α* = 0.1. Here, smm+IMIX controlled the FDR for each data type individually, and the univariate and multivariate IMIX models controlled an across-data-type FDR at 0.1. Multivariate IMIX and univariate IMIX discovered more genes than smm+IMIX. In particular, 1 533 genes that were not identified by the smm+IMIX method were discovered by both integrative spatial methods. All three proposed methods discovered 715 genes of interest. Multivariate IMIX identified 443 field-effect genes that were not found using the other two methods, while univariate IMIX discovered 349 such genes. On the basis of the simulation study results, next we concentrated on the univariate IMIX results. We visualized the field-effect genes and filtered the genes for which negative correlations between the gene expression data and methylation data were preserved. There were 1 512 genes that met these filtering criteria. The unsupervised clustering of the log2ratio values of genes based on methylation value and the visualizations of both methylation and gene expression levels are shown in Figure 6. The identified genes showed field effect across all NU/LGIN, HGIN, and UC groups. The hypermethylated and hypomethylated genes showed constant log2ratio changes across groups compared with the controls. The gene expression values were negatively correlated with methylation values, with the same patterns across all groups. Figure 7 highlights an extracted part of the genes mapped to chromosome 18. This visualization corresponds to the spatial locations of bladder mucosa in the 3D maps. We observed similar patterns to those described for the 2D heatmaps (Figure 6).

**Figure 6:**
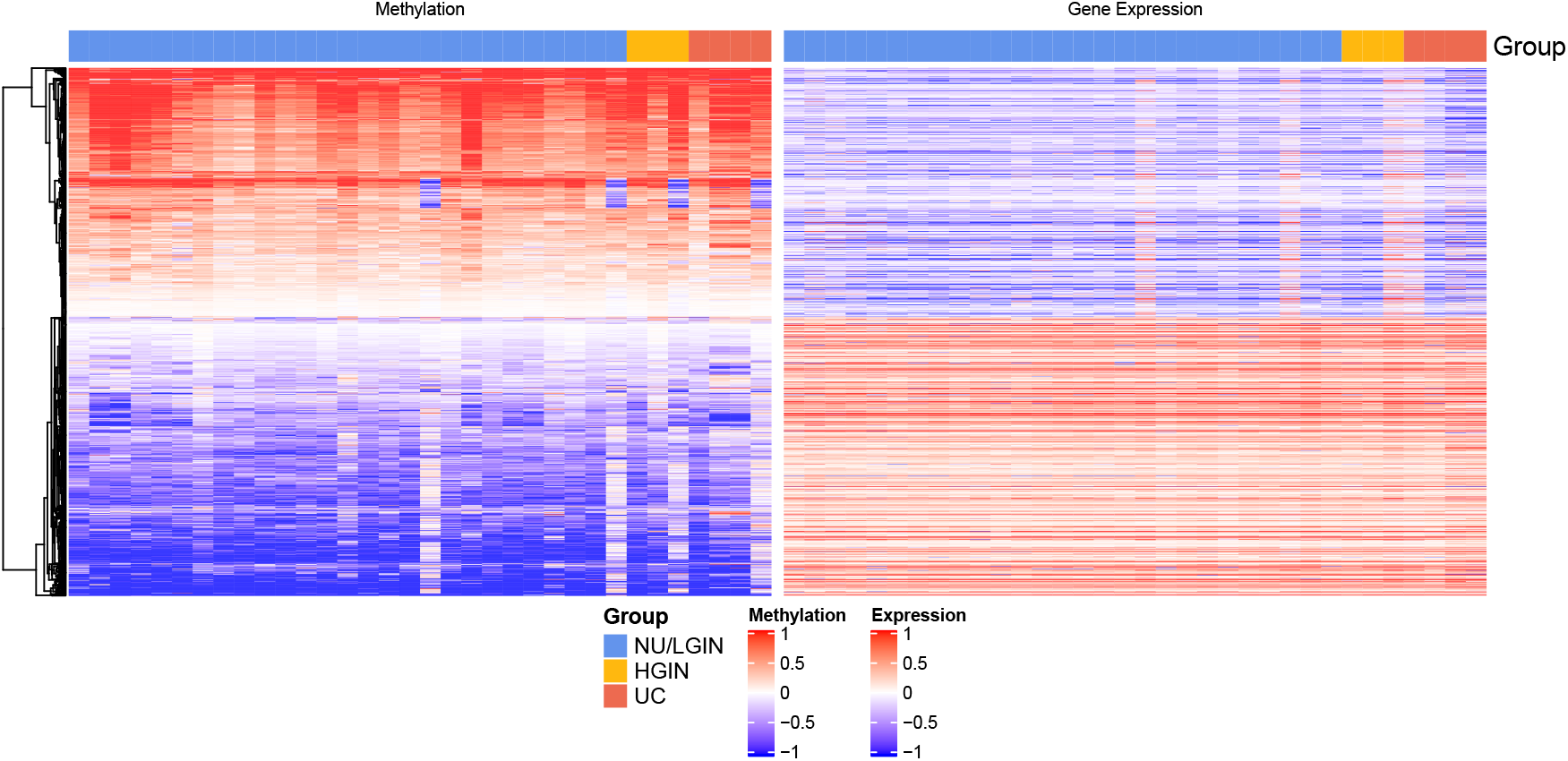
Visualization of the log2ratio of the methylation and gene expression values compared with the controls of the filtered field effect genes (*N* = 1 512) identified by univariate IMIX at FDR=0.1.

**Figure 7:**
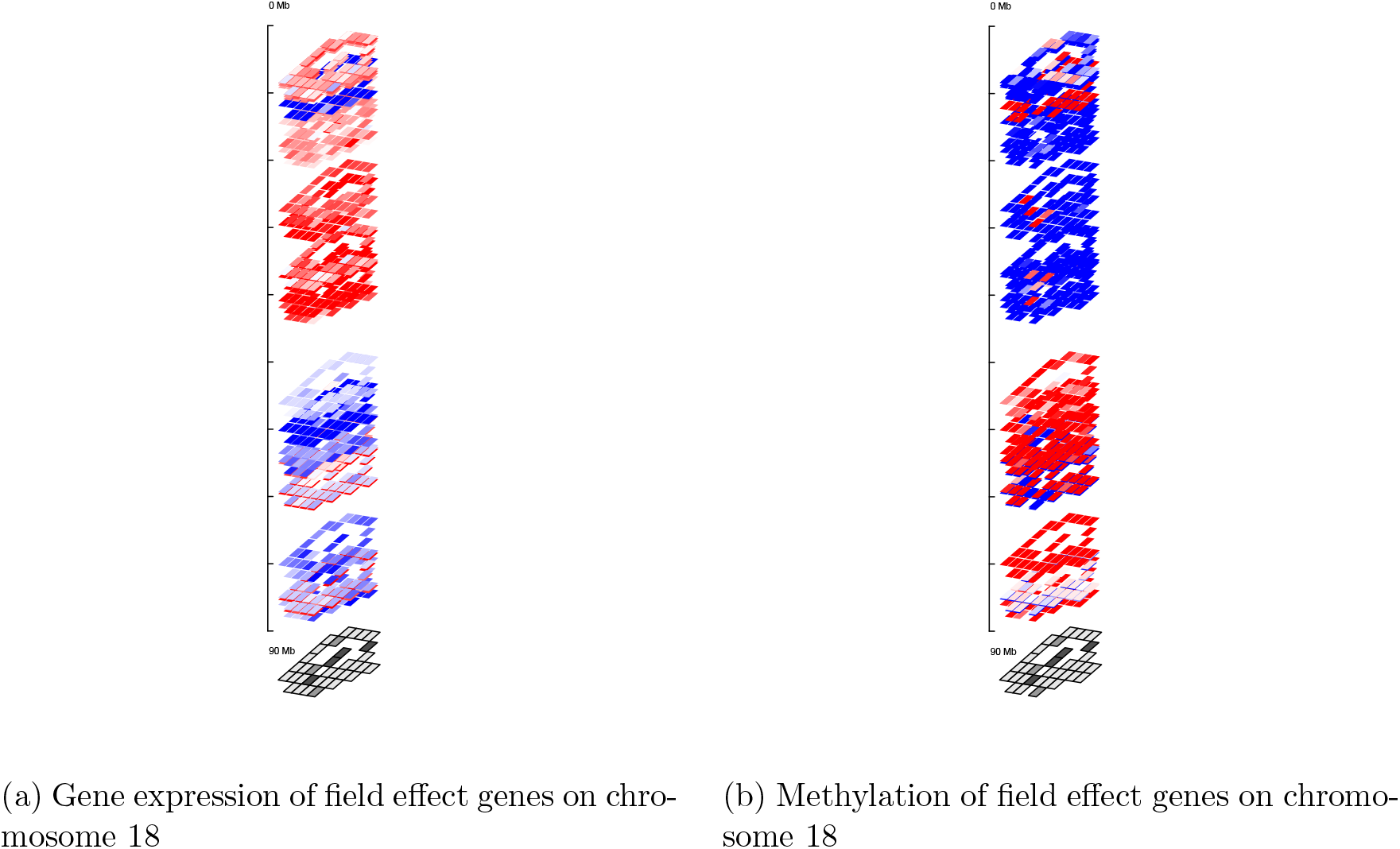
3D pattern of the log2ratio of (a) gene expression and (b) methylation values compared with the controls of the field effect genes on chromosome 18 (*N* = 26) identified by univariate IMIX at FDR=0.1.

The identified genes control important cell functions in the development of bladder cancer. We performed pathway analysis using IPA on the field-effect genes discovered by the three methods with FDR controlled at *α* = 0.01 of gene expression and methylation data. The 78 significant pathways with Benjamini-Yekutieli (BY) FDR control at 0.05 detected by univariate spatial IMIX are visualized together with the pathways detected by the other two methods (Figure S4). There was no significant pathway after BY FDR control using the separate analysis smm+IMIX. The discovered pathways can be categorized into three groups: immunity and inflammation, signal transduction-differentiation, and carcinogenesis. Among which, we found p53 signaling, PPAR, regulation of the Epithelial-Mesenchymal Transition (EMT), and tumor microenvironment pathways. These are closely related with tumor initiation, growth, and invasion.

Our integrative method has been shown to have more power in detecting field-effect genes in both data types by simultaneously taking into account the complicated dependence structures between two data types and their spatial correlation structures. Furthermore, other methods such as SPARK and SpatialDE are not suitable for this type of application, in which the genes do not show any variations related to the spatial locations, but rather an overall same-direction differential expression/methylation pattern across all spatial locations. Our analysis using SPARK is described in the supplementary materials Section 2.2. It was able to discover genes showing variations based on spatial locations but was not able to discover specific disease-group-label-related genes, such as NU/LGIN, HGIN, or UC disease grades.

## 4 Discussion

Spatial IMIX is a multivariate mixture model framework for spatially correlated multi-omics data integration. It can identify differentially expressed/methylated genes associated with prespecified patterns of sample subtypes. It is a model-based method that incorporates the spatial correlation structures between samples in a geographically resolved area by applying a spatial mixed model, which improves the power to detect genes in certain sample subtypes within each genomic dataset. Spatial IMIX additionally considers the dependence structures between different genomic datasets by assuming a multivariate Gaussian mixture distribution of the *z*-scores (transformed from *P* -values) from the spatial mixed model of individual-level data. We showed that spatial IMIX had a lower misclassification rate and better controlled FDR than did existing methods such as linear regression and spatial linear mixed models in the simulation study.

Our method, which considers spatially correlated data of multiple omics data types and sample subtypes, is an extension of our previously developed IMIX method (Wang and Wei, 2020) for multi-omics data integration for association analysis. Spatial IMIX retains the advantages of IMIX, including great computational efficiency from its use of the EM algorithm for whole-genome wide analysis, its improved model fitting of mixture models by using model selection based on BIC values, and the error-control properties of multiple hypothesis testing by using an adaptive procedure to control the FDR among different sample subtypes across the omics data types. Additionally, the fact that the data integration step only considers the summary statistics enables us to use independent or partially overlapping sample locations, as illustrated in the whole-organ mapping of bladder cancer application (section 3.2), where the measured samples with gene expression data and methylation data were not completely spatially aligned. The two proposed models, univariate IMIX and multivariate IMIX, both showed good power in the simulation study. Specifically, univariate IMIX, a parsimonious and flexible version of spatial IMIX, can be easily extended to multiple types of data and sample subtypes. This is an improvement of IMIX, which only considered two or three types of data.

Spatial IMIX could be further applied to spatial transcriptomics data with more complicated sample subtypes and larger sample sizes. Spatial transcriptomics is a cutting-edge technology that measures spatially resolved gene expression at high throughput. Several novel methods have been proposed to identify genes with spatial patterns of expression variation, such as spatialDE (Svensson et al., 2018) and SPARK (Sun et al., 2020). These methods could be used for unsupervised clustering to identify unknown or non-prespecified patterns in a geographical area for each gene. Spatial IMIX is different from previously proposed methods, which mainly aimed to find spatially variable genes without considering the location sample subtypes. Our method, in contrast, focuses on fixed effect estimations given spatially resolved data with stringent FDR control for the identified genes associated with certain prespecified patterns of sample subtypes. In particular, spatialDE and SPARK are not suitable for the type of applications described in section 3.2 where the genes did not vary on the basis of their spatial locations, but instead in terms of differential expression or methylation in the same overall direction across all spatial locations. With the ongoing development of spatial transcriptomics and other large-scale multi-omics data technologies, our work has a wide range of potential applications that could provide novel biological insights into different problems involving spatially resolved data, such as the incorporation of information about tissue makeup and sample subtypes.

We implemented the data integration in the mixture model framework in a way that accounts for the correlation structures of two omics data types through the mixing proportions. Further investigations of the complicated correlation structures of the variance-covariance matrix estimations could be explored and compared with our current parsimonious univariate and multivariate spatial IMIX models. We leave this potential extension for future research. Our method relaxed the conditional independence assumptions for multiple data types in the presence of spatial correlation through a two-step procedure using univariate mixed models as the first step to consider the spatial correlation and multivariate mixture model as the second step to consider the multi-omics data correlations. Nonetheless, the development of multivariate spatial mixed models for the data application presented in this paper could be an interesting future direction.

