## Supplemental materials for "Spatial IMIX: A Mixture Model Approach to Spatially Correlated Multi-Omics Data Integration"

### Supplementary Materials for "Spatial IMIX: A Mixture Model Approach to Spatially Correlated Multi-Omics Data Integration"

Ziqiao Wang[1], Bogdan Czerniak[2,*], and Peng Wei[3,*]

[1]Department of Biostatistics, Bloomberg School of Public Health, Johns Hopkins University, Baltimore, MD, USA
[2]Department of Pathology, The University of Texas MD Anderson Cancer Center, Houston, TX, USA
[3]Department of Biostatistics, The University of Texas MD Anderson Cancer Center, Houston, TX, USA

#### 1 Simulation Studies

##### 1.1 Simulation Study: Type I Error and Power Comparisons

The joint hypothesis testing whether the gene shows differentially expressed/methylated patterns in any group (Table 1 & 3) is:

$$H_0 \; : \; \beta_0 = \beta_1 = \beta_2 = 0;$$

$$H_1 \; : \; \text{At least one is not 0.}$$

We also tested the type I error and power for each fixed effect estimate (Table 2 & 4). To implement the testings, we used R packages spaMM (Rousset and Ferdy, 2014) and nlme (Pinheiro et al., 2021) and both performed well using the likelihood ratio test (LRT). Given the computational time consideration, we adapted nlme R package for spatial mixed modeling in our method.

| Spatial Var | lm:Ftest | lm:Chisq | smm:Wald | smm+LRT |
|---|---|---|---|---|
| 0.1 | 0.077 | 0.094 | 0.099 | 0.046 |
| 0.5 | 0.165 | 0.193 | 0.166 | 0.062 |
| 1 | 0.228 | 0.258 | 0.194 | 0.069 |
| 2 | 0.292 | 0.328 | 0.207 | 0.071 |

Table 1: Typer I error of fixed effects comparisons at 0.05, this is the joint testing of all three fixed effect estimates using linear regression (lm) based on F test and Wald test, and spatial mixed model (smm) using Wald test and LRT.

| Spatial Var | lm:LG | lm:HG | lm:UC | smm+Ftest:LG | smm+Ftest:HG | smm+Ftest:UC | smm+LRT:LG | smm+LRT:HG | smm+LRT:UC |
|---|---|---|---|---|---|---|---|---|---|
| 0.1 | 0.066 | 0.059 | 0.060 | 0.057 | 0.055 | 0.057 | 0.004 | 0.060 | 0.063 |
| 0.5 | 0.123 | 0.080 | 0.089 | 0.093 | 0.068 | 0.075 | 0.011 | 0.070 | 0.078 |
| 1 | 0.163 | 0.094 | 0.107 | 0.111 | 0.072 | 0.083 | 0.016 | 0.073 | 0.084 |
| 2 | 0.199 | 0.111 | 0.127 | 0.123 | 0.073 | 0.089 | 0.020 | 0.075 | 0.086 |

Table 2: Typer I error of fixed effects comparisons at 0.05, this is the separate testing of each of three fixed effect estimates using linear regression (lm), spatial mixed model (smm) using F test, and spatial mixed model using LRT.

| Fixed Effect | Spatial Var | lm:Ftest | lm:Chisq | smm:Wald | smm+LRT |
|---|---|---|---|---|---|
| 1 | 0.1 | 1.000 | 1.000 | 1.000 | 0.924 |
| 2 | 0.1 | 1.000 | 1.000 | 1.000 | 1.000 |
| 3 | 0.1 | 1.000 | 1.000 | 1.000 | 1.000 |
| 2 | 0.5 | 1.000 | 1.000 | 1.000 | 0.999 |
| 2 | 1 | 1.000 | 1.000 | 1.000 | 0.994 |
| 2 | 2 | 1.000 | 1.000 | 1.000 | 0.958 |

Table 3: Power of fixed effects comparisons at 0.05, this is the joint testing of all three fixed effect estimates using linear regression (lm) based on F test and Wald test, and spatial mixed model (smm) using Wald test and LRT.

| Fixed Effect | Spatial Var | lm:LG | lm:HG | lm:UC | smm+Ftest:LG | smm+Ftest:HG | smm+Ftest:UC | smm+LRT:LG | smm+LRT:HG | smm+LRT:UC |
|---|---|---|---|---|---|---|---|---|---|---|
| 1 | 0.1 | 0.994 | 0.674 | 0.675 | 0.990 | 0.660 | 0.656 | 0.862 | 0.675 | 0.666 |
| 2 | 0.1 | 1.000 | 0.997 | 0.997 | 1.000 | 0.996 | 0.996 | 0.999 | 0.996 | 0.995 |
| 3 | 0.1 | 1.000 | 1.000 | 1.000 | 1.000 | 1.000 | 1.000 | 1.000 | 1.000 | 1.000 |
| 2 | 0.5 | 1.000 | 0.976 | 0.974 | 1.000 | 0.973 | 0.969 | 0.988 | 0.975 | 0.969 |
| 2 | 1 | 1.000 | 0.929 | 0.925 | 0.999 | 0.923 | 0.910 | 0.931 | 0.926 | 0.911 |
| 2 | 2 | 0.996 | 0.817 | 0.805 | 0.981 | 0.802 | 0.776 | 0.754 | 0.809 | 0.781 |

Table 4: Power of fixed effects comparisons at 0.05, this is the separate testing of each of three fixed effect estimates using linear regression (lm), spatial mixed model (smm) using F test, and spatial mixed model using LRT.

#### 1.2 Simulation Study: Integrative Analysis

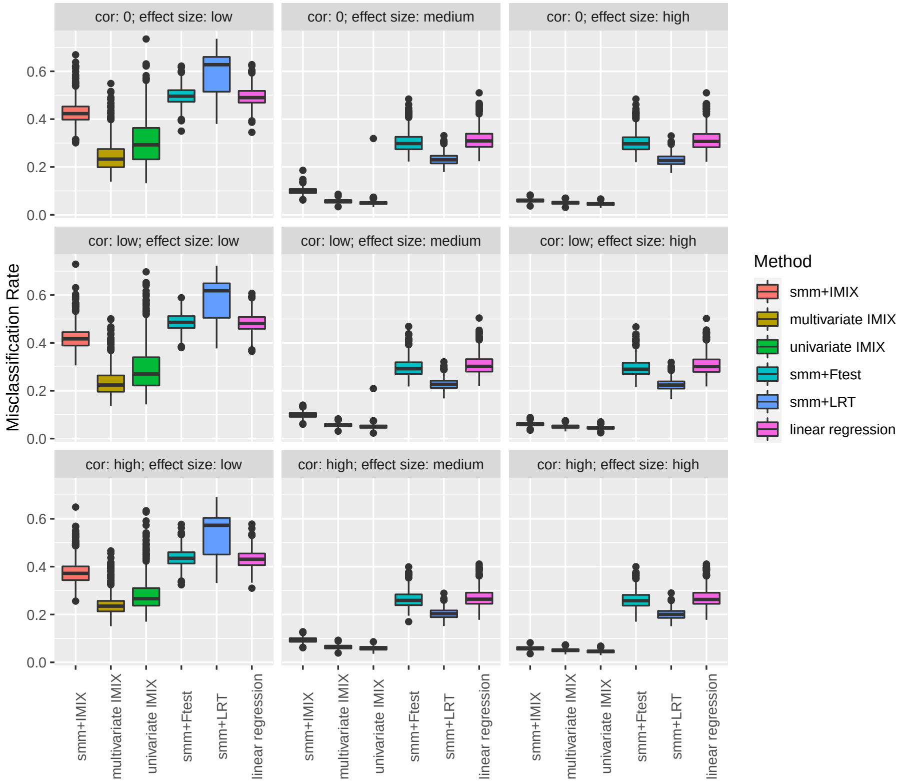


Figure 1: Simulation study results for two omics data types comparing spatial mixed model (smm) with IMIX extension, multivariate spatial IMIX, univariate spatial IMIX, smm with F test, smm with LER, and linear regression using different data correlation and effect size settings (Scenario 1-3).

#### 1.3 Simulation Study: Model Selection

| | Model Selection | Effect Size | | |
|---|---|---|---|---|
| | | Low | Medium | High |
| Multivariate IMIX | Before | 0.225824074 | 0.059104497 | 0.050149471 |
| | After | 0.198121693 | 0.056972222 | 0.049208995 |
| Univariate IMIX | Before | 0.272166667 | 0.05330291 | 0.045437831 |
| | After | 0.221944444 | 0.052002646 | 0.045328042 |

Table 5: Comparison of misclassification rate of fitted four-component mixture model after BIC model selection and 64-component mixture model before model selection at FDR=0.2 for simulation study.

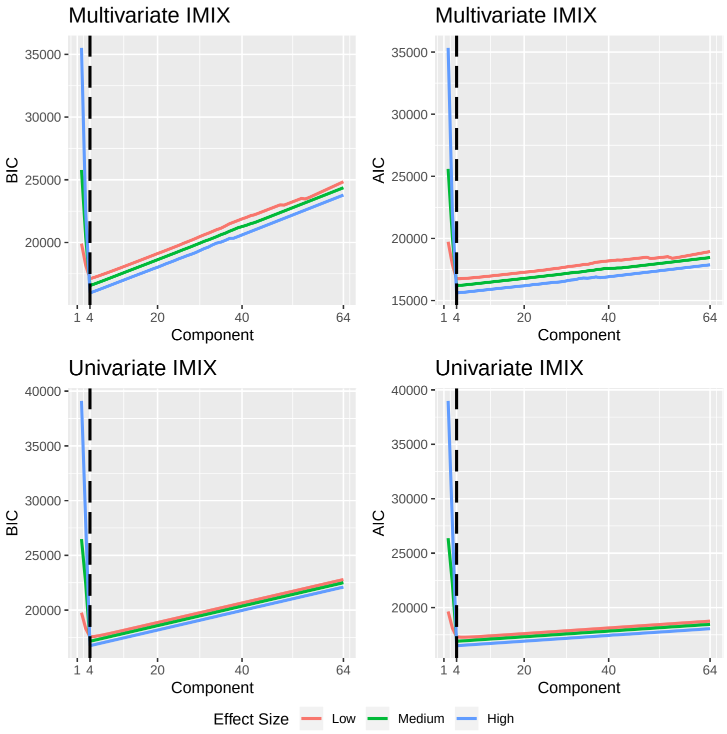


Figure 2: Average AIC and BIC values of 1 000 simulation studies for each component based on different effect size settings.

### 2 Real Data Applications to The Whole Organ Mapping of Bladder Cancer

#### 2.1 Data Preprocessing and Quality Control

**Methylation Data Preprocessing and Gene Level Summary.** Methylation data were measured on the Illumina Infinium HumanMethylation450 (450K) BeadChip array at over 480,000 sites. The CpG sites with missing values in more than 10% of the total samples were filtered out. We excluded all the probes on the sex chromosomes, the probes of target polymorphic CpGs that overlaps with known SNPs (Fortin et al., 2017), and the probes that are cross-reactive (Chen et al., 2013).We preprocessed and normalized the raw IDAT files for each sample using the minfi R package (version 1.30.0). Gene-level methylation data were summarized based on Spearman correlations between the CpG sites within each gene and its expression level based on the results of an external dataset, the TCGA bladder cancer tissue samples. We represent the methylation level for each gene as of the single probe that has the most negative correlation between methylation and expression in the 391 TCGA samples by interrogating either the 1st exon, 5'UTR, or up to 1500bp upstream from the transcription start site (TSS). The data set in total contains 14 744 genes with 27 LGIN, 3 HGIN, 4 UC samples in map 19, and 8 normal controls for downstream analysis.

**RNA Sequencing Data Preprocessing.** The RNA integrity and RIN number was assessed using a 2100 Bioanalyzer (Agilent). RNA concentration was determined using RiboGreen quantification (Quant-iT RiboGreen RNA Assay Kit from Invitrogen). RNA samples meeting a quantity threshold of 1 µg with the RNA integrity number (RIN) $\geq 7$ were analyzed by the Advanced Technology Genomics (ATCG) Core. There were 34 mucosal samples from map 19 and five sex-matched normal control urothelial suspensions, which were prepared from the ureters of nephrectomy specimens that were free of urothelial neoplasia. RNA sequencing alignment was done using STAR/2.7.2b with BAM files as the output. We used DESeq2 for data normalization and batch effect corrections for protein-coding only genes. The data set in total contained 18 692 genes with 27 LGIN, 3 HGIN, 4 UC samples in map 19 and 5 normal controls for downstream analysis.

For methylation data:

| Compnent in IMIX | 1 | 2 | 3 | 4 | 5 | 6 | 7 | 8 |
|---|---|---|---|---|---|---|---|---|
| FDR=0.05 | 9186 | 5327 | 3 | 0 | 0 | 0 | 166 | 0 |
| FDR=0.1 | 5201 | 9227 | 5 | 0 | 0 | 0 | 249 | 0 |
| FDR=0.2 | 3892 | 10187 | 7 | 210 | 0 | 3 | 380 | 3 |

Table 6: Significant genes in each component of IMIX with across-data-type FDR for exponential spatial structure mixed model

| | No effect | Field effect | LG-,HG+,UC+ | LG-,HG-,UC+ | Others |
|---|---|---|---|---|---|
| FDR=0.05 | 9186 | 5327 | 166 | 0 | 65 |
| FDR=0.1 | 5201 | 9227 | 249 | 0 | 67 |
| FDR=0.2 | 3892 | 10190 | 380 | 210 | 72 |

Table 7: Significant genes in each group by IMIX fitting and LRT p-values with across-data-type FDR for exponential spatial structure mixed model

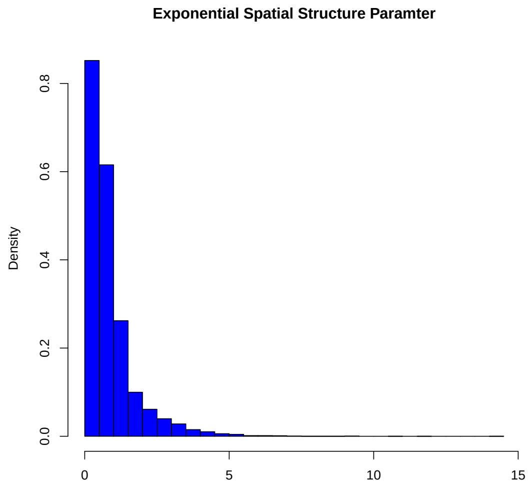


Figure 3: Parameter distribution of the exponential spatial structure

For RNAseq data map 19:

| Compnent in IMIX | 1 | 2 | 3 | 4 | 5 | 6 | 7 | 8 |
|---|---|---|---|---|---|---|---|---|
| FDR=0.05 | 17621 | 975 | 1 | 7 | 0 | 0 | 0 | 0 |
| FDR=0.1 | 16543 | 1990 | 5 | 50 | 0 | 0 | 0 | 16 |
| FDR=0.2 | 14509 | 3539 | 27 | 459 | 0 | 0 | 0 | 70 |

Table 8: Significant genes in each component of IMIX with across-data-type FDR for exponential spatial structure mixed model-RNAseq map19

| | No effect | Field effect | LG-,HG+,UC+ | LG-,HG-,UC+ | Others |
|---|---|---|---|---|---|
| FDR=0.05 | 17621 | 975 | 0 | 7 | 89 |
| FDR=0.1 | 16543 | 1991 | 0 | 50 | 108 |
| FDR=0.2 | 14509 | 3548 | 0 | 459 | 176 |

Table 9: Significant genes in each group by IMIX fitting and LRT p-values with across-data-type FDR for exponential spatial structure mixed model-RNAseq map19

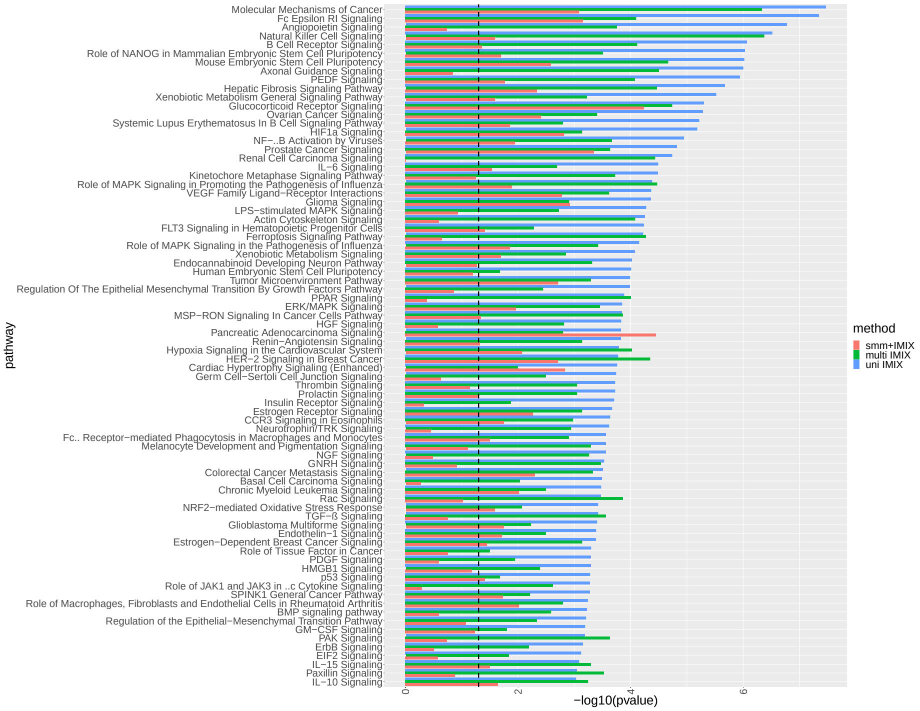


Figure 4: The canonical pathways after Benjamini-Yekutieli FDR control at 0.05 identified by the Ingenuity Pathway Analysis (IPA) on the field-effect genes identified by univariate spatial IMIX with adaptive FDR controlled at $\alpha = 0.1$. Dashed line is -log10(0.05).

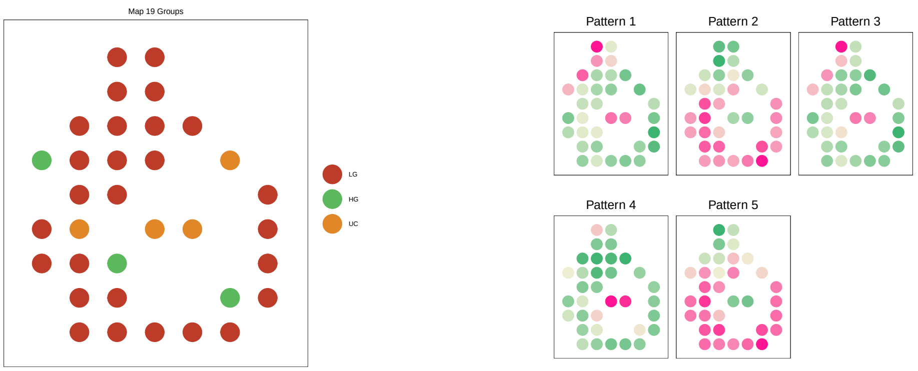


(a) (b)

Figure 5: SPARK results on the whole organ mapping of bladder cancer. (a) Disease subtype visualization. (b) Average methylation level of discovered genes grouped by hierarchical clustering.

#### 2.2 SPARK Analysis

We analyzed the methylation data of whole organ mapping data of bladder cancer using the Gaussian version of SPARK (Sun et al., 2020) and used hierarchical agglomerative clustering algorithm to distinguish the patterns. As suggested in their manuscript, we set the two optional parameters in the R function to be Euclidean distance and Ward's criterion. We clustered the discovered genes with spatial variation into 5 groups, the visualization of the mean values for genes in each pattern is shown in Figure 5. As shown here, SPARK was not able to discover field-effect genes (genes without spatial variation) but only genes that showed a different spatial pattern in the tissue area. In addition, it was not possible to have statistical significance levels for the genes grouped in each pattern with respect to the disease subtype labels using this method.
